# Complex dietary-polysaccharide modulates gut immune function and microbiota, and promotes protection from autoimmune diabetes

**DOI:** 10.1101/492637

**Authors:** Radhika Gudi, Nicolas Perez, Benjamin M. Johnson, M.Hanief Sofi, Robert Brown, Songhua Quan, Subha Karumuthil-Melethil, Chenthamarakshan Vasu

**Affiliations:** Department of Microbiology and Immunology, College of Medicine, Medical University of South Carolina, Charleston, SC-29425; University of Illinois at Chicago, Chicago, IL; Department of Radiation Oncology, Northwestern University Feinberg School of Medicine, Chicago, IL; Department of Neurology, University of North Carolina, Chapel Hill, NC

**Keywords:** Yeast β-glucan, gut mucosa, innate immunity, autoimmunity, Type 1 diabetes, Gut microbiota, immune regulation, Beta-glucan, dietary polysaccharide, mucosal immunity, Dectin-1, immune modulation/tolerance/suppression

## Abstract

Since the dietary supplement and prebiotic value of β-glucan-rich products have been widely recognized and the dietary approaches for modulating autoimmunity have been increasingly explored, we assessed the impact of oral administration of high-pure yeast β-glucan (YBG) on gut immune function, microbiota and type 1 diabetes (T1D) using mouse models. Oral administration of this non-digestible complex polysaccharide caused a Dectin-1-dependent immune response involving increased expression of IL10, retinaldehyde dehydrogenase (Raldh) and pro-inflammatory cytokines in the gut mucosa. YBG-exposed intestinal DCs induced/expanded primarily Foxp3+, IL10+ and IL17+ T cells, ex vivo. Importantly, prolonged oral administration of low-dose YBG at pre-diabetic stage suppressed insulitis and significantly delayed the T1D incidence in non-obese diabetic (NOD) mice. Further, prolonged treatment with YBG showed increased Foxp3+ T cell frequencies, and a significant change in the gut microbiota, particularly an increase in the abundance of Bacteroidetes and a decrease in the Firmicute members. Oral administration of YBG, together with Raldh-substrate and β-cell antigen, resulted in a better protection of NOD mice from T1D. These observations suggest that YBG not only has a prebiotic property, but also has an oral tolerogenic-adjuvant-like effect, and these features could be exploited for modulating autoimmunity in T1D.

## INTRODUCTION

Beta-glucans (β-glucans; BGs) are non-digestible complex polysaccharides, commonly found in plants, cereals, bacteria, yeast, and fungi e.g. mushrooms. They can potentiate the function of immune system and improve the antimicrobial activity of innate immune cells ^1, 2^. BGs can activate both innate and adaptive immunity and hence, have been investigated as an effective systemic adjuvant in immunotherapies for cancer and infections ^1, 2^. Many of the BG-containing preparations have been promoted as daily dietary supplements to enhance immune and metabolic functions ^3^, and are suggested to have prebiotic properties. Our recent report showed that systemic injection of low amounts of highly purified BG (purity >98%) from Baker’s yeast (*Saccharomyces cerevisiae*) (YBG) modulates autoimmune progression and prevents type 1 diabetes (T1D) in non-obese diabetic (NOD) mice ^4^, suggesting it has previously unknown immune regulatory properties. Structurally, YBG contains β-1,3-linked D-glucose molecules (β-1,3-D-glucan) with β-1,6-linked side chains and it is recognized by innate immune cells, primarily through a pathogen recognition receptor (PRR) called Dectin-1^5, 6^. While clinical studies using YBG preparations have shown beneficial effects ^7-10^, a clinical trial failed to detect changes in either cytokine production or microbicidal activity of leukocytes upon oral administration of YBG ^11^. Overall, since most of these published studies on biological effects of BGs have administered BGs systemically, and/or used partially purified and very poorly defined products, the degree and nature of immune modulation induced by purified BGs and the impact of oral consumption of BGs on autoimmunity are largely unknown.

Dietary approaches have increasingly been considered for modulating immune function, and preventing autoimmunity and other clinical conditions. Therefore, in this study, we examined the gut immune and microbiota modulatory properties of orally administered high-pure YBG (1-3,1-6-β-glucan), and the impact on organ specific autoimmunity in T1D using NOD mice. We observed that oral administration of YBG resulted in the expression of higher levels of immune regulatory and pro-inflammatory cytokines, and non-cytokine factors such as vitamin A metabolizing enzyme, retinaldehyde dehydrogenase (Raldh), in the gut mucosa. This mixed innate immune response also contributed to an increase in the regulatory T cell (Treg) frequency. YBG recipient mice also showed a significantly higher abundance in the microbial communities of Bacteroidetes and Verrucomicrobia phyla, to which many polysaccharide-fermenting bacteria belong, and decrease in Firmicute members, compared to untreated controls. While oral administration of YBG was sufficient to significantly delay the hyperglycemia in NOD mice, co-administration of YBG in an antigen specific manner [β-cell-Antigen (β-cell-Ag)] and retinol further potentiated the diabetes protection effect in these mice. Our observations suggest that a combination of YBG-induced innate immune response in the gut mucosa and altered microbiota contributes to the suppression of autoimmune progression in NOD mice, and these features can be exploited for preventing T1D in at-risk subjects by using dietary supplement based immune modulators.

## Materials and Methods

### Mice

C57BL/6, NOD/LtJ, OT-II, NOD-BDC2.5, and NOD-*Scid* mice were purchased from the Jackson laboratory. C57BL6-Dectin-1 deficient (Dectin-1^-/-^) mice ^12^ were provided by Dr. Iwakura, University of Tokyo, Japan. Foxp3-GFP-knockin (Foxp3-GFP) mice were provided by Dr. Vijay Kuchroo (Harvard Medical School, MA). Breeding colonies of these strains were established and maintained in the pathogen-free facility of University of Illinois at Chicago (UIC) or Medical University of South Carolina (MUSC). NOD-Foxp3-GFP mice were generated by backcrossing B6-Foxp3-GFP-ki mice to NOD background for 12 generations. To generate NOD-BDC-Foxp3-GFP mice, NOD-BDC2.5 mice were crossed with NOD-Foxp3-GFP mice. To detect hyperglycemia, glucose levels in blood collected from the tail vein of wild-type NOD/Ltj and NOD-*Scid* mice were tested using the Ascensia Micro-fill blood glucose test strips and an Ascensia Contour blood glucose meter (Bayer, USA). Mice with glucose levels >250 mg/dl for two consecutive weeks were considered diabetic. All animal studies were approved by the animal care and use committees at UIC and MUSC.

### β-cell-Ag, peptides and other Reagents

Immunodominant β-cell antigen peptides [viz., 1. Insulin B (9-23), 2. GAD65 (206-220), 3. GAD65 (524-543), 4. IA-2beta (755-777), 5. IGRP (123-145); 6. BDC2.5 TCR reactive peptide (YVRPLWVRME; referred to as BDC-peptide), and 7. OVA (323-339) peptides (ISQAVHAAHAEINEAGR; referred to as OT-II peptide) were custom synthesized (Genscript Inc) and used in this study. Peptides 1-5 were pooled at an equal molar ratio and used as β-cell-Ag as described earlier ^13-15^. Highly purified YBG (glucan from Baker’s yeast, *S. cerevisiae*), ≥98% pure, was purchased from Sigma-Aldrich and the purification method has been described before ^16^. This agent was tested for specific activity using thioglycolate-activated macrophages as described before ^4, 17, 18^. L-tryptophan, Retinol, PMA, ionomycin, Brefeldin A, and purified and conjugated antibodies were purchased from Sigma-Aldrich, BD Biosciences, eBioscience, Invitrogen, R&D Systems, and Biolegend Laboratories. Magnetic bead-based total and CD4+ T cells, and CD11c+ dendritic cell isolation kits, as well as magnetic-bead linked secondary antibodies (Miltenyi Biotec, Invitrogen and StemCell technologies) were used for enriching or depleting cell populations. In some experiments, intestinal epithelial cells were depleted by Percoll gradient centrifugation or using A33 antibody (Santa Cruz Biotechnology, CA) and anti-rabbit IgG-magnetic beads. ELISA and magnetic bead-based suspension array reagents/kits were purchased from R&D Systems, BD Biosciences, Invitrogen, Bio-Rad, Affymetrix-eBioscience and/or Millipore.

### Treatments

Six to eight-week old C57BL/6 and NOD-BDC-Foxp3-GFP, 10-week-old pre-diabetic (glucose levels <100 mg/dl) female NOD/Ltj mice were given YBG suspension in saline (250 μg/mouse/day) for different durations by oral gavage. Control mice were given saline. In some experiments, mice were given a higher dose of YBG (2000 μg/mouse/day). In some experiments, mice were also given synthetic peptides (5 μg/mouse/day) or Retinol (0.5 μg /mouse/day) by oral gavage. For depleting gut microbiota, C57BL/6 mice were given a broad-spectrum antibiotic cocktail (ampicillin (1 g/l), vancomycin (0.5 g/l), neomycin (1 g/l), and metronidazole (1 g/l) - containing drinking water for up to 30 days.

### Dendritic cells and ex vivo assays

Dendritic cells were isolated from the ileum of control and YBG-fed mice. Intestinal tissues were cleaned thoroughly by flushing with PBS-containing antibiotics, cut into small pieces, incubated in collagenase (0.5 mg/ml) and DNAse (50 μg/ml) containing RPMI medium for 1 h, passed through 70 um filter, and washed using RPMI medium with 10% FBS. Percoll gradient was generated by layering 100%, 40%, and 25% Percoll in which intestinal cells were suspended, and centrifuged for 15 min at 2000g. Cells at the interphase of 100% and 40% Percoll layers were collected as immune cell-rich fraction for dendritic cell (DC) isolation. Intestinal CD11c^+^ DCs were enriched from immune cell fraction using magnetic bead-linked anti-CD11c Ab reagent. In some experiments, cells were stained using fluorochrome-labeled CD11c and CD103 antibodies, and CD11c+CD103+ and CD11c+CD103-populations were enriched using a MoFlo high speed cell sorter. In some experiments, transcript levels of various factors in these cells and cytokines spontaneously secreted by them, upon ex vivo culture for 48h without additional stimulation, were determined. In antigen presentation assays, purified T cells (1x10^5^ cells/well) were incubated with antigenic peptide-pulsed DCs (2x10^4^ DCs/well). In some assays, these cultures were supplemented with retinol and TGFβ1. For some assays, *in vitro* stimulated or freshly isolated T cells were re-stimulated using PMA (50 ng/ml) and ionomycin (500 ng/ml) in the presence of Brefeldin A (1 μg/ml) for 4h before staining for intracellular cytokines. In some assays, spleen, mesenteric lymph node (MLN) and pancreatic lymph node (PnLN) cells (2 x 10^5^ cells/well) from treated and control mice were stimulated with anti-CD3 Ab (1 µg/ml) for 24 h or β-cell-Ag (5 μg/ml) for 48 h and spent media were tested for cytokines.

### Adoptive T cell transfer experiment

Pancreatic LN (PnLN) cell suspension of control and YBG fed mice were transferred into 6-8-wk-old NOD*-Scid* mice (i.v.; 1 x 10^6^ cells/mouse); glucose levels were tested every week.

### Histochemical analysis of pancreatic tissues

Pancreata were fixed in 10% formaldehyde, 5-µm paraffin sections were cut, and stained with hematoxylin and eosin (H&E). Stained sections were analyzed using a grading system in which 0 = no evidence of immune cell infiltration, 1 = peri-islet infiltration (<5%), 2= 5-25% islet infiltration, 3 = 25–50% islet infiltration, and 4 = >50% islet infiltration and completely destroyed islets, as described in our earlier studies ^13-15, 19^.

### 16S rRNA gene targeted sequencing and bacterial community profiling

Total DNA was prepared from the fecal pellets for bacterial community profiling as detailed in our previous reports ^20, 21^. DNA in the samples was amplified by PCR using 16S rRNA gene-targeted primer sets to assess the bacterial levels. V3-V4 region of 16S rRNA gene sequencing was performed using Illumina MiSeq platform at MUSC genomic center. The sequencing reads were fed in to the Metagenomics application of BaseSpace (Illumina) for performing taxonomic classification of 16S rRNA targeted amplicon reads using an Illumina-curated version of the GreenGenes taxonomic database which provided raw classification output at multiple taxonomic levels. The sequences were also fed into QIIME open reference operational taxonomic units (OUT) picking pipeline ^22^ using QIIME pre-processing application of BaseSpace. The OTUs were compiled to different taxonomical levels based upon the percentage identity to GreenGenes reference sequences (i.e. >97% identity) and the percentage values of sequences within each sample that map to respective taxonomical levels were calculated. The OTUs were also normalized and used for metagenomes prediction of Kyoto Encyclopedia of Genes and Genomes (KEGG) orthologs employing PICRUSt as described before ^23-26^. The predictions were summarized to multiple levels and functional categories were compared among control and YBG fed groups using the statistical Analysis of Metagenomic Profile Package (STAMP) as described before ^24^.

### Statistical analysis

Mean, SD, and statistical significance (*p-value*) were calculated, graphic visualizations including heatmaps were made using Excel (Microsoft), Prism (GraphPad), Morpheus (Broad Institute), and STAMP ^24^ applications. One or two-tailed *t*-test was employed, unless specified, for values from *in vitro* and *ex vivo* assays where two groups were compared. Log-rank analysis was performed to compare T1D (hyperglycemia) incidences of two groups. Fisher’s exact test was used for comparing number of immune cell infiltrated islets in test vs. control groups. All the statistical analyses for microbial sequences were done employing two-tailed *t*-test and the *p*-values were corrected from multiple tests using Benjamini and Hochberg approach in STAMP. A *p*-value of ≤0.05 was considered significant.

## Results

### Oral administration of YBG results in Dectin-1 dependent gut immune activation

Previous studies have suggested that partially purified BG and BG-containing agents can activate the gut immune system ^27-29^. In this study, WT and Dectin-1-/-mice were given high-pure (>98%), YBG (high-dose: 2000 μg/mouse/day or low-dose: 250 μg/mouse/day) for 3 days, and overall immune phenotype of the small intestine was determined. Fig. 1A shows that small intestine of WT mice, but not Dectin-1-/-mice, expressed significantly higher levels of cytokine and non-cytokine factors compared to that of untreated mice. YBG-induced gut innate immune response in WT mice involved higher expression of mRNA both immune regulatory and pro-inflammatory factors including IL10, Raldh1A2, TNFα, IL6 and IL1β. Similar analysis of distal colon showed relatively higher expression of a few cytokine and chemokine factors in YBG treated mice (not significant statistically; not shown). These results suggest the ability of purified YBG to induce Dectin-1 dependent immune response in the gut mucosa. Of note, short-treatment with low-dose YBG induced only modestly higher, not statistically significant, levels of some cytokine and non-cytokine factors in the small intestine (Supplemental fig. 1).

**FIGURE 1:**
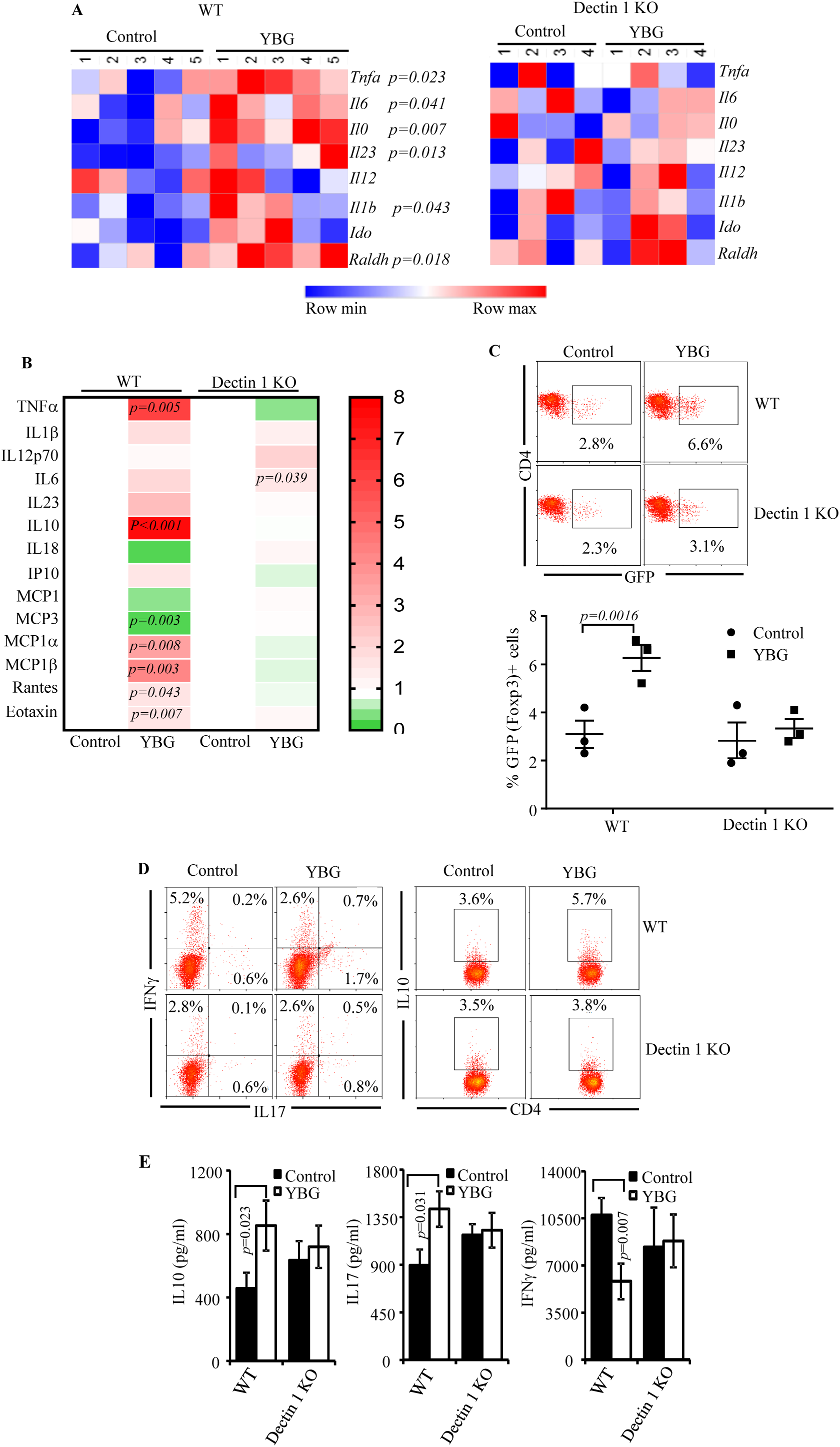
Oral administration of YBG causes Dectin-1-dependent immune activation in the gut mucosa. **A)** Eight-week-old WT and Dectin-1^-/-^ B6 mice were given YBG suspension (2000 μg/mouse/day) by oral gavage for 3 consecutive days and euthanized after 24 h to harvest the intestine. cDNA prepared from the distal ileum region was subjected to qPCR and the expression levels of cytokines and non-cytokine factors were compared. Expression levels relative to β-actin expression were plotted as a heatmap with Morpheus application. n= 4-5 mice/group and the assay was performed in triplicate for each sample. Similar analysis of distal colon did not show significant differences (not shown). **B)** WT and Dectin-1^-/-^ B6 mice were treated and euthanized as described above, small intestinal CD11c+ DCs were MACS-sorted from IEC-depleted (using A33 antibody), enriched intestinal immune cell fraction. Cells were cultured in triplicate for 48h (2x10^5^/well in 96-well plate) and supernatants were tested for the levels of spontaneously secreted cytokines by suspension bead array. Mean values of untreated WT and Dectin-1^-/-^ mouse cells were considered as baseline value of 1 for calculating fold cytokine secretion by cells from YBG treated mice. n= 3 pools of cells (2-3 mice/pool). Heatmap was generated using GraphPad Prism. **C**) Freshly isolated DCs were pulsed with OT-II peptide and cultured (2x10^4^ cells/well) with OT-II-Foxp3-GFP mouse splenic T cells (1x10^5^ cells/well) for 4 days and examined for CD4+ cells with GFP (Foxp3) expression. Representative FACS scatter plots (left) and mean±SD values of three independent experiments in triplicate (right) are shown. **D)** DCs and OT-II mouse T cells were cultured similarly, restimulated using PMA and ionomycin for 4 h and stained for intracellular cytokines. Representative FACS scatter plots of three independent experiments are shown. **E)** Supernatants collected from the cultures described for panel C were tested by ELISA for cytokine levels. Mean±SD values of three independent parallel experiments in triplicate are shown. These experiments were repeated at least twice. Statistical significance was calculated by *t-*test.

### Intestinal DCs (IDCs) from YBG fed mice produce higher amounts of cytokines and skew T cell response ex vivo

Since intestinal immune cells, DCs in particular, are known to encounter dietary and microbial agents in the gut lumen ^30-32^, in vivo response of IDCs to short-term treatment with high-dose YBG was examined. As shown in Fig. 1B, IDCs from WT mice, but not Dectin-1-/-mice, that received YBG orally showed higher secretion of pro-and anti-inflammatory cytokine and chemokine factors compared to respective untreated controls. Since YBG exposure caused activation of IDCs in vivo, these cells were tested for their ex vivo ability to promote T cell activation and differentiation upon antigen presentation. IDCs from YBG fed mice could induce significantly higher frequencies of Foxp3-, IL10-and IL17A-expressing/producing T cells, as compared to their counterparts from untreated mice (Fig. 1C-E). On the other hand, compared to control mouse IDCs, these cells from YBG fed mice induced lower IFNγ response in T cells. These observations show that YBG exposure activates IDCs and enhances their immune modulatory/regulatory properties.

### CD103^+^ IDCs, but not CD103^-^ IDCs, of YBG treated mice induce/expand Foxp3+ T cells

Since CD103+ cells are the key IDC population that expresses the tolerogenic enzyme Raldh1A2 and other immune regulatory factors as well as directly encounters gut luminal content ^30, 33, 34^, and oral administration of YBG caused higher expression of mRNA for these factors in the intestine, CD103^+^ and CD103^-^ IDCs of YBG fed mice were sorted and tested for the expression levels of mRNA for key cytokine and non-cytokine factors. As shown in Fig. 2A, CD103^+^ IDCs from high-dose YBG fed mice expressed higher levels of mRNA for Raldh1A2 and IL-10 compared to their counterparts from control mice while CD103^-^ IDCs from these YBG fed mice expressed relatively higher levels of mRNA for IL6. Ex vivo antigen presentation assay revealed that while CD103^+^ IDCs from YBG fed mice induced significantly higher frequencies of IL10-producing T cells, CD103^-^ IDCs from these mice induced IL17-producing T cells (Fig. 2B). Further, CD103^+^ and CD103^-^ IDCs from YBG fed mice induced lower and higher IFNγ production respectively, albeit not significant statistically, compared to their counterparts from control mice (not shown). Importantly, CD103^+^ IDCs from YBG fed mice showed superior ability to induce/expand Foxp3^+^ T cells, in the absence or presence of Raldh-substrate (retinol) and TGFβ1, compared to their counterparts from control mice (Fig. 2C). These observations suggest that YBG exposure activates both CD103^+^ and CD103^-^ IDCs and promotes distinct T cell responses.

**FIGURE 2:**
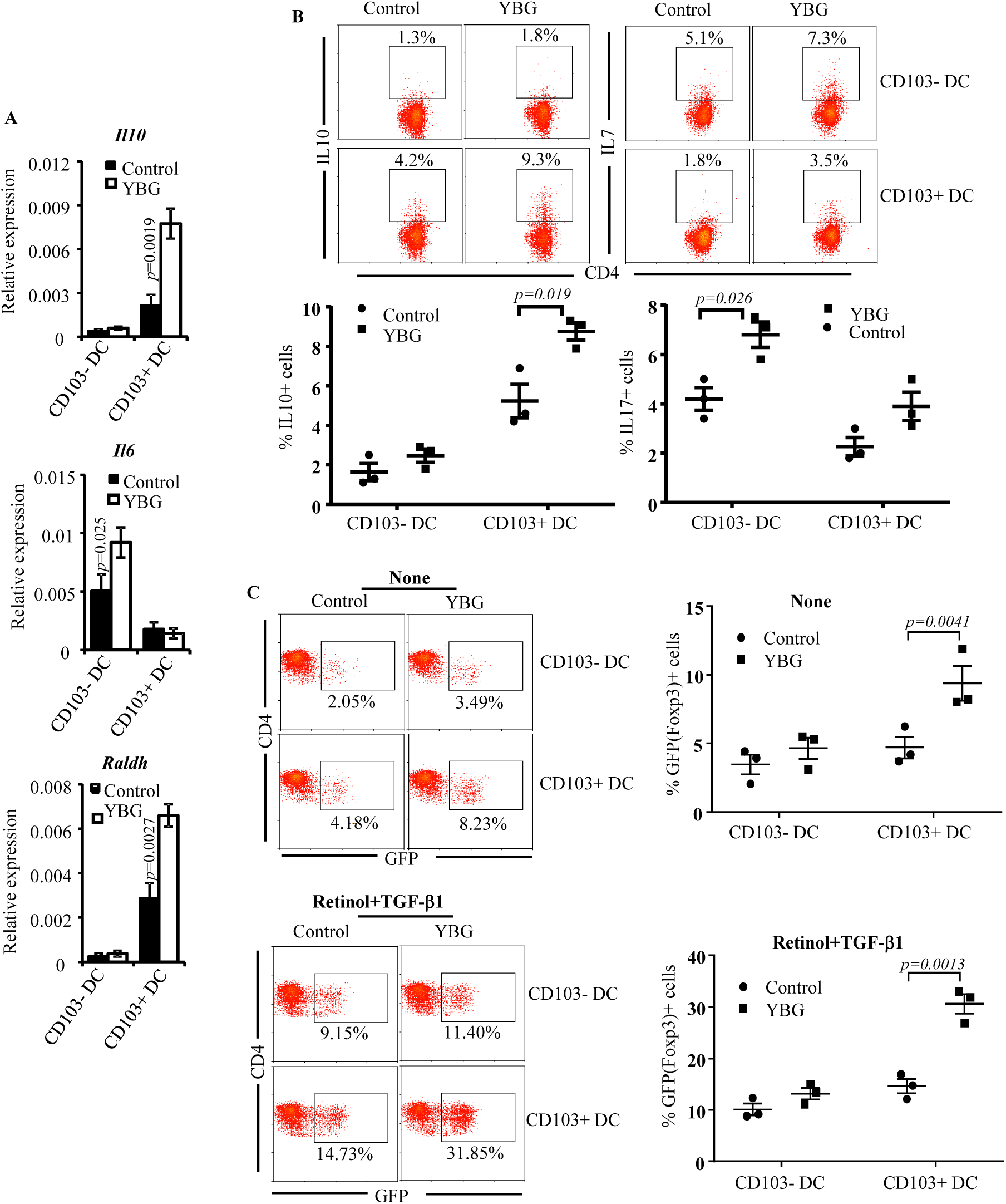
CD103+ IDCs from YBG treated mice induce T cells with regulatory phenotype ex vivo. WT B6 mice were treated and euthanized as described for Figure 1, small intestinal CD11c+CD103-and CD11c+CD103+ DCs were FACS-sorted from IEC-depleted, enriched intestinal immune cell fraction. **A**) Freshly isolated cells were subjected to qPCR assay. Expression levels relative to β-actin expression are shown. n= 3 pools of cells (4-5 mice/pool). **B**) OT-II peptide pulsed DCs and OT-II T cells were cultured and processed for detection of intracellular cytokines as described for Fig. 1. Representative FACS scatter plots (upper panel) and mean±SD values of three independent experiments conducted in triplicate (lower panel) are shown. **C)** DCs were cultured similarly with OT-II-Foxp3-GFP T cells in the absence or presence of retinol and examined for CD4+ T cells with GFP expression. Representative FACS scatter plots (left) and mean±SD values of three independent experiments in triplicate (right) are shown. These experiments were repeated at least thrice. Statistical significance was calculated by *t*-test.

### Prolonged oral administration of YBG protects pre-diabetic NOD mice from T1D

For above described short-term studies, which were conducted using B6 mice due to the availability of Dectin-1 KO mice in that background, high dose YBG (2000 μg/mouse/day) was used to reliably assess the Dectin-1 dependent and overall in vivo effect of orally administered YBG. However, since oral administration of YBG resulted in upregulation of both immune regulatory and pro-inflammatory factors, for experiments involving long-term treatment, clinically relevant low-dose YBG (250 μg/mouse/day) is desired as a precautionary measure to minimize the likelihood of intestinal inflammation upon prolonged treatment. Modulation of gut immune function upon short-term treatment with high-dose YBG prompted studies to determine the impact of low-dose YBG consumption on gut immune phenotype and pancreatic autoimmunity using a spontaneous model of T1D, which was employed in our previous study to determine the impact of systemic injection of YBG^4^. As observed in Supplemental fig. 2A, gut mucosa of YBG treated NOD mice showed cytokine and Raldh1A2 mRNA expressions similar to that of B6 mice (shown in Fig. 1 and Supplemental fig. 2B). Importantly, pre-diabetic NOD mice that were given low-dose YBG for 30 consecutive days showed significantly delayed hyperglycemia (Fig. 3A). This treatment prevented the disease in more than 40% of the mice for at least 18 weeks post-treatment, whereas 100% of the control mice developed diabetes within this period. One set of mice were euthanized, 7 days post-treatment and pancreatic tissues were examined for insulitis severity. The pancreatic islets of YBG treated mice showed significantly less severe immune cell infiltration and insulitis, compared to untreated mice (Fig. 3B). While about 50% of the islets in mice that received β-glucan showed insulitis grade ≤2, at least 60% of the islets in untreated euglycemic mice showed insulitis severity grade >2. These results suggest that immune modulation induced by orally administered YBG leads to curbed immune cell infiltration and β-cell destruction, resulting in delayed onset of hyperglycemia in NOD mice.

**FIGURE 3:**
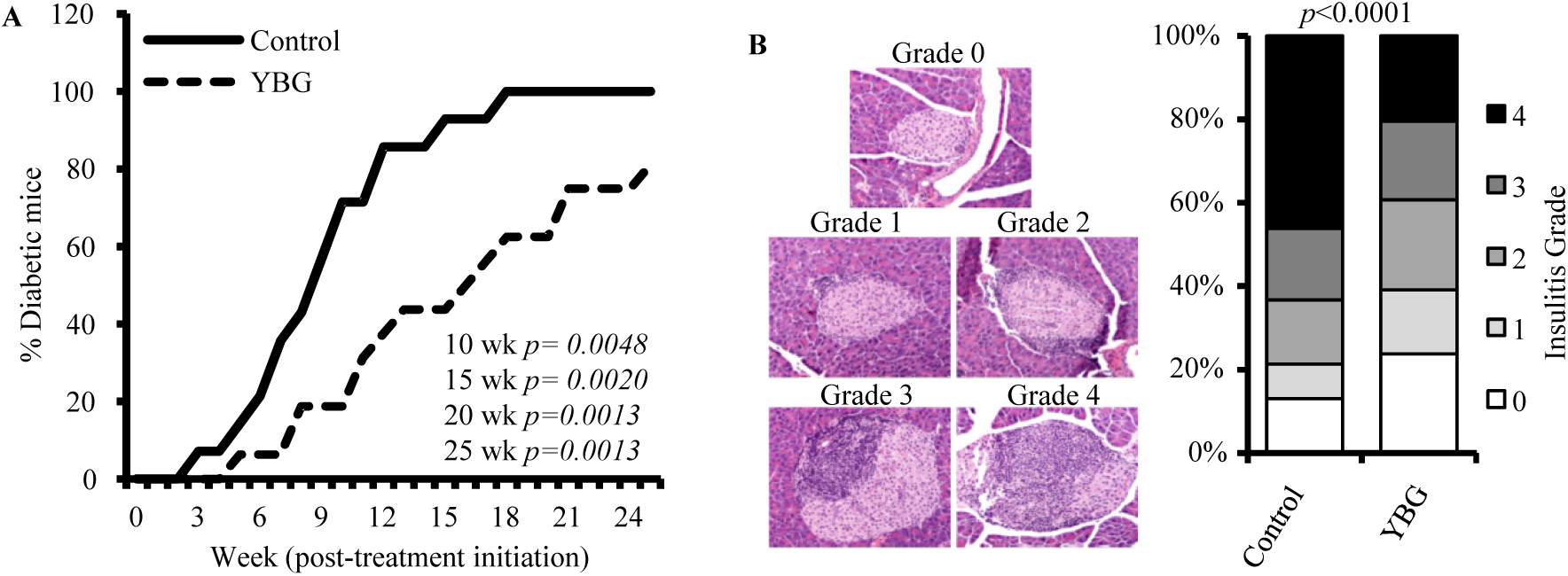
Oral administration of low-dose YBG resulted in delayed hyperglycemia in NOD mice. Female NOD mice were given YBG suspension (250 μg/mouse/day) or saline by oral gavage for 30 consecutive days starting at 10 weeks of age. **A)** One set of mice (n=14 for control and 16 for YBG groups) were tested for blood glucose levels, weekly for up to 25 weeks post-treatment initiation, to detect hyperglycemia/diabetes. **B)** One set of mice (5/group) were euthanized one-week post-treatment and H&E stained pancreatic sections were examined for insulitis. Representative islets with insulitis grades (**left**) and overall percentage of islets with different insulitis grades (**right**) are shown. Statistical significance was calculated by Log-rank test (for panel A) and Fisher’s exact test (for panel B).

### Prolonged oral administration of YBG results in modulation of T cell phenotype and function in the pancreatic LNs

Since YBG-fed mice showed less severe insulitis and significant delay in hyperglycemia, T cells from these mice were examined for their phenotypic and functional properties in comparison with their counterparts from control mice. Foxp3+ T cell frequencies were profoundly higher in the intestinal mucosa and the PnLNs of YBG fed mice compared to control mice (Fig. 4A). Importantly, both β-cell Ag (Fig. 4B) and PMA/ionomycin (not shown) stimulations ex vivo, followed by FACS analysis, showed that IL10+, IL17+, and IL4+ T cell frequencies were significantly higher and IFNγ+ T cell frequency is relatively lower in the PnLN of YBG fed mice compared to controls. Further, levels of these cytokines secreted by PnLN cells from YBG fed and control mice in response to β-cell-Ag challenge ex vivo showed similar trends (Fig. 4C). To determine if YBG treatment impacted the function of immune cells, diabetogenic property of PnLN cells was examined by adoptively transferring these cells from YBG treated and control mice into NOD-*Scid* mice. Fig. 4D shows that NOD-*Scid* mice that received immune cells from YBG-treated mice developed hyperglycemia relatively slower compared to control cell recipients. Further, pancreatic tissues of NOD-*Scid* mice that received cells from YBG treated mice showed significantly lower insulitis severity compared to control mouse cell recipients (Fig. 4E). Overall, these observations suggest that prolonged YBG consumption results in immune modulation not only in the gut mucosa, but also in the systemic compartment including pancreatic microenvironment, thereby causing the suppression of insulitis in NOD mice.

**FIGURE 4:**
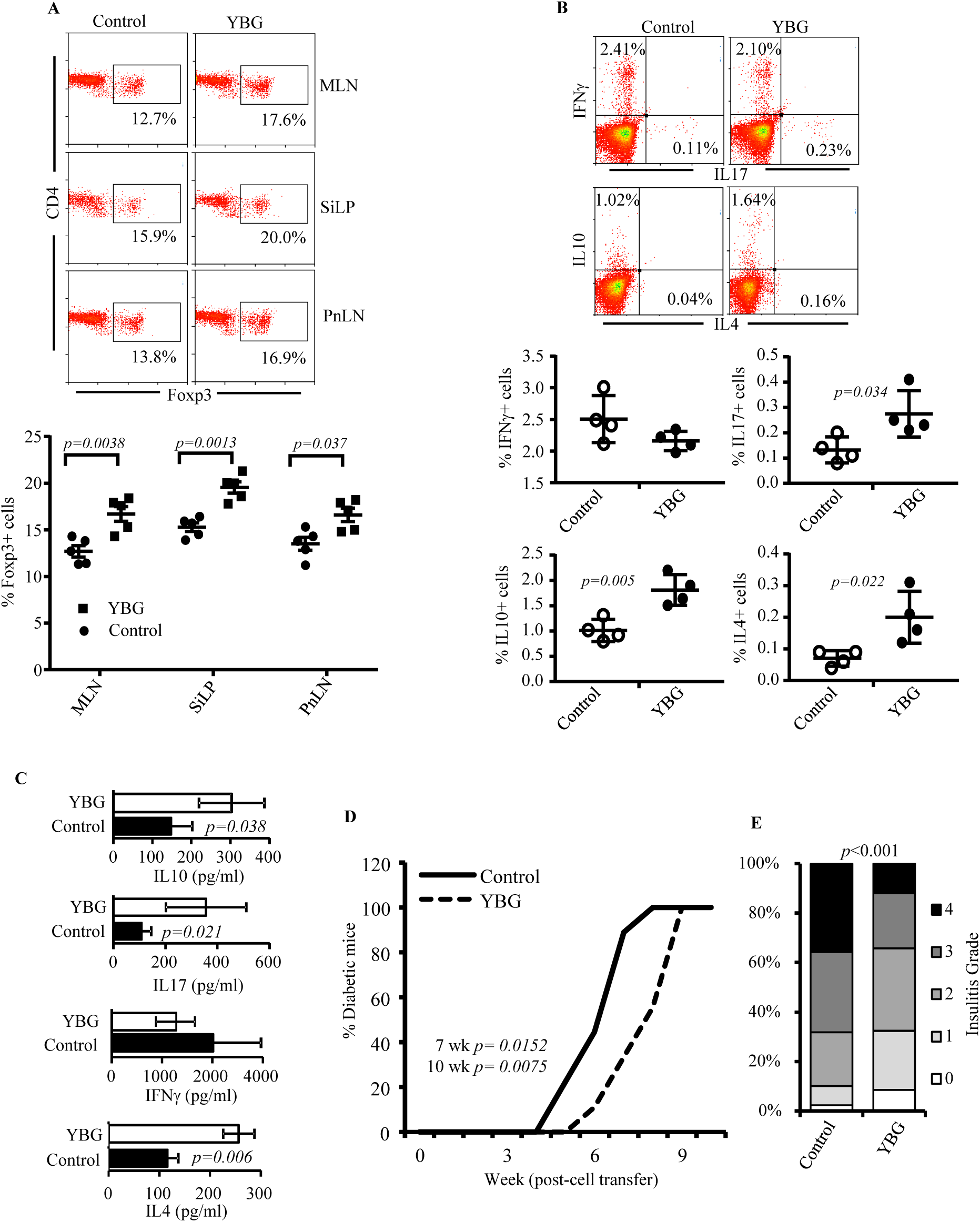
YBG treatment caused modulation of T cell function in the systemic compartment and suppression of autoimmunity. Female NOD mice were given low-dose YBG by oral gavage for 30 consecutive days starting at 10 weeks of age as described for Fig. 5 and the mice were euthanized one-week post-treatment. **A)** MLN, SiLP and PnLN cells were stained for Foxp3 expression and analyzed by FACS. **B)** PnLN cells from these mice were stimulated using β-cell-Ag for 24h and examined for intracellular cytokine levels by FACS. Representative FACS plots (upper panels) and mean±SD values of 4 to 5 mice (lower panel) are shown for A and B. **C)** PnLN cells were stimulated using β-cell-Ag for 48 h and the supernatants were examined for cytokines. **D)** PnLN cells from at least 5 treated and 5 control mice were pooled separately and injected i.v. into NOD-*Scid* mice (1x10^6^ cells/mouse; 9 recipients/group) and monitored for blood glucose levels to detect diabetes. **E)** One set of mice (3/group) were euthanized 3 weeks post-cell transfer and H&E stained pancreatic sections were examined for insulitis. Overall percentage of islets with different insulitis grades are shown. Statistical significance was calculated by *t*-test (for panels A-C), log-rank test for up to 7 and 10 weeks (for panel D), and Fisher’s exact test (for panel E).

### Prolonged oral administration of low-dose YBG results in changes in the gut microbiota

Complex non-digestible dietary polysaccharides are thought to be degraded / fermented by colonic bacteria and affect the overall gut microbiota composition and immune function^35-37^, and it could impact the autoimmune outcomes. Therefore, we determined the impact of prolonged oral treatment of NOD mice with YBG on gut microbiota. The mice were given YBG by oral gavage every day for 30 days, fresh fecal pellets from treated and control mice were collected, and subjected to 16S rRNA targeted gene sequencing to determine the profiles of microbial communities. Compilation of OTUs to different taxonomical levels showed that the abundances of major phyla were significantly different in YBG fed mice, compared to control mice (Fig. 5A). YBG treated mice showed significantly higher abundance of Bacteroidetes and Verrucomicrobia phyla members and profoundly lower abundance of Firmicutes and Proteobacteria in their fecal samples compared to controls. Examination of OTUs at genus level revealed that the abundances of many major microbial communities are significantly different in YBG treated and control mice (Fig. 5B). Overall, major shift in favor of microbial communities belonging to Bacteroidetes and Verrucomicrobia, which include many polysaccharide-degrading bacterial communities, over Firmicutes occurred in YBG treated mouse gut. For OTU-based prediction of metabolic functions of gut microbes that may be associated with complex dietary polysaccharide degradation and metabolite production, PICRUSt application was employed. Predicted major pathways that are overrepresented in YBG treated mice, based on 16S rRNA gene identity, include carbohydrate metabolism, biosynthesis of secondary metabolites, energy metabolism, cofactor and vitamin metabolism and xenobiotics biodegradation and metabolism (Fig. 5C). Overrepresentation of these pathways in gut microbiota of YBG fed mice suggests that oral consumption of this dietary polysaccharide can cause selective enrichment of carbohydrate fermenting/consuming-and metabolite-producing bacteria in the gut.

**FIGURE 5:**
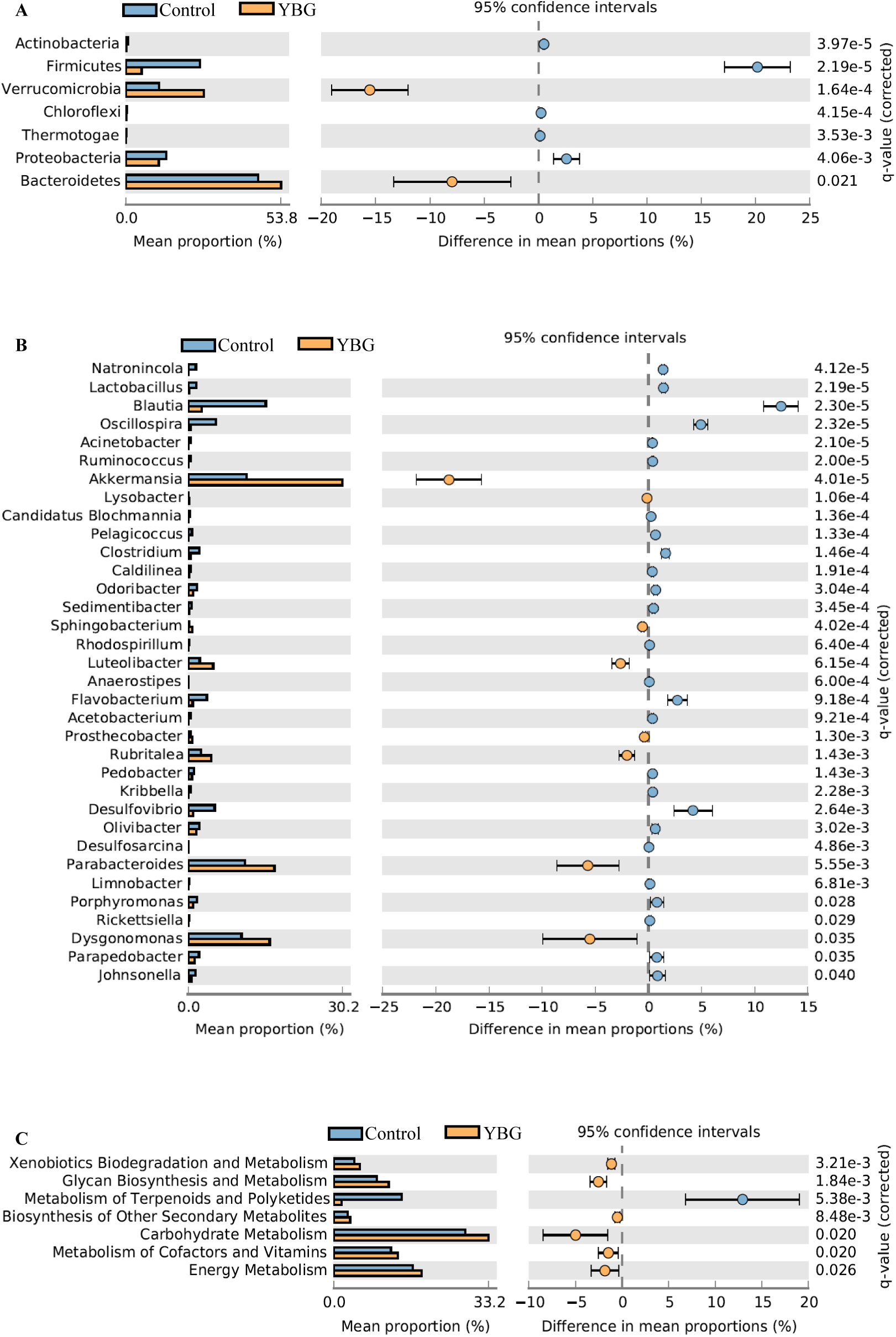
Prolonged oral administration of low-dose YBG results in changes in the composition of gut microbiome in NOD mice. Ten week old pre-diabetic NOD mice were given saline or YBG suspension (250 μg/mouse/day) for 30 consecutive days by oral gavage as described for Fig. 3, and fresh fecal pellets from individual animals (4/group) were collected post-treatment, DNA samples prepared from these fecal pellets were subjected to 16S rRNA gene (V3V4) -targeted sequencing using the Illumina MiSeq platform. The OTUs that were compiled to different taxonomical levels based upon the percentage identity to reference sequences (i.e. >97% identity) and the percentage values of sequences within each sample that map to specific phylum and genus were calculated. **A**) Relative abundances of 16S rRNA gene sequences in fecal samples of individual mice before and after treatment at phyla level are shown. **B**) Mean relative abundances of sequences representing the major microbial communities (>1 % of total sequences) in YBG treated group at genus level are shown. **C**) Normalized OTU-biom table was used for predicting gene functional categories using PICRUSt application and selected level 2 categories of KEGG pathway are shown. Statistical analyses were done employing *t*-test and the *p*-values were corrected from multiple tests using Benjamini and Hochberg approach in STAMP.

### YBG treatment associated protection from T1D is, in part, gut microbe dependent

Since YBG treatment of NOD mice resulted in significant changes in the composition of gut microbiota, the influence of microbiota on YBG treatment associated protection of NOD mice from T1D was examined. Pre-diabetic mice were given a cocktail of broad spectrum antibiotics in drinking water to deplete gut microbiota (Supplemental fig. 3) and treated concurrently with YBG. Microbiota depleted control mice showed relatively rapid progression of insulitis and higher incidence of hyperglycemia, compared to untreated control mice with intact microbiota (Fig. 6A). On the other hand, microbiota-depleted mice showed only a modest protection from hyperglycemia upon YBG treatment compared to mice that received YBG alone. Correspondingly, overall severity of insulitis in microbiota depleted mice that received YBG was found to be considerably higher than that observed in YBG treated mice with intact microbiota, compared to microbiota depleted control mice (Fig. 6B). These observations suggest the possibility that both gut immune modulation induced directly by YBG and indirectly by altered gut microbiota contribute to the protection of NOD mice from T1D upon oral consumption of YBG. These observations prompted examination of Treg frequencies in the PnLN of YBG treated NOD mice without and with microbial depletion. Fig. 6C shows, similar to Fig. 4A, significantly higher frequencies of Foxp3+ T cells in the PnLN of YBG treated mice with intact gut microbota compared to their control counterparts. Importantly, while depletion of gut bacteria alone caused substantial reduction in Foxp3+ T cell frequencies in both control and YBG recipient mice compared to their counterparts with intact microbiota, PnLN from microbiota-depleted YBG recipient NOD mice showed significantly higher frequencies of Foxp3+ T cells compared to microbiota depleted controls. These observations show that while gut microbes appear to contribute to the maintenance of overall Treg function, oral consumption of YBG has a direct impact on immune activation and T cell phenotype.

**FIGURE 6:**
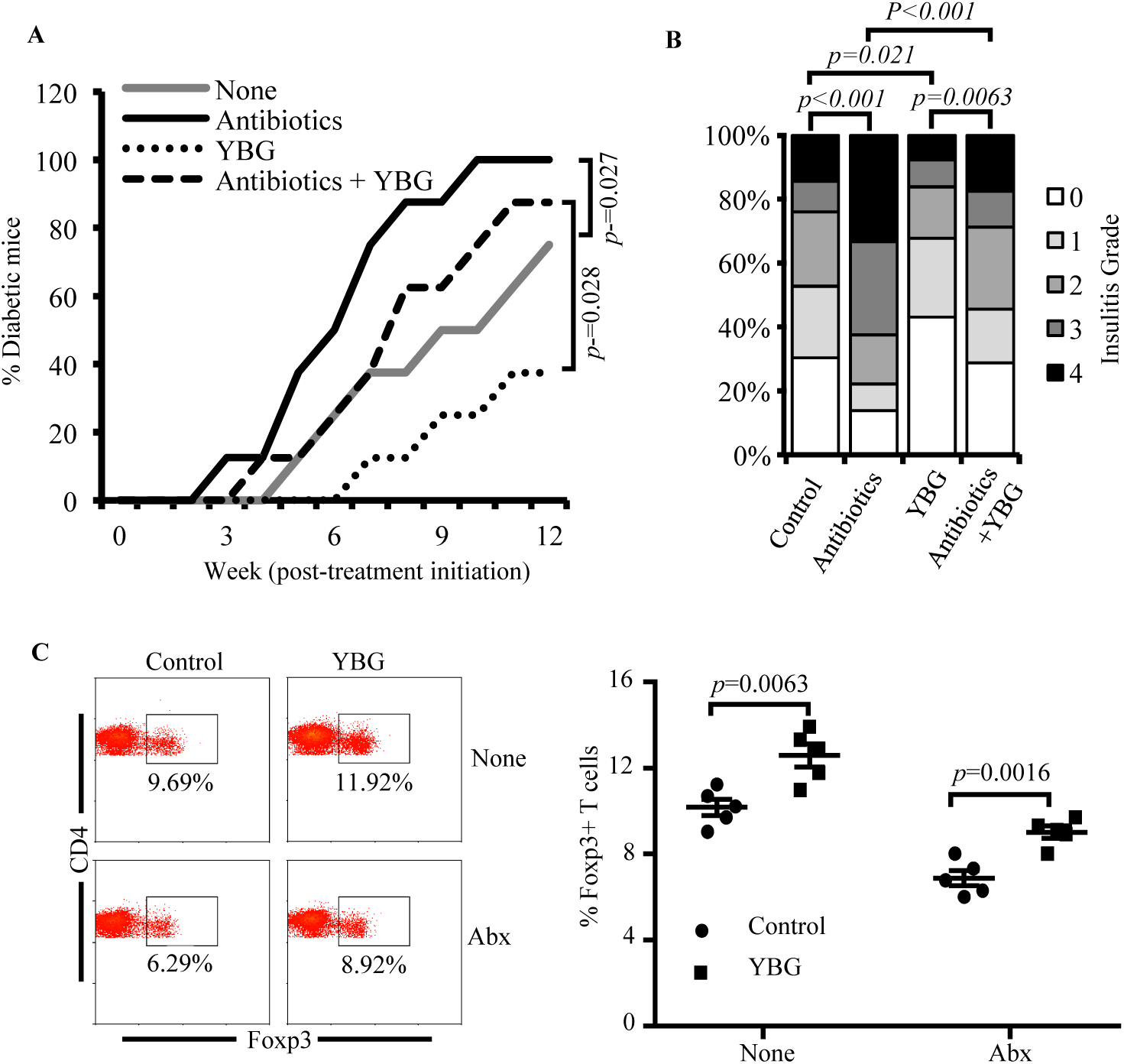
YBG treatment associated protection of NOD mice from T1D is partially gut microbe dependent. Female NOD mice were given YBG for 30 consecutive days starting at 10 weeks of age as described above for Fig. 5, and maintained on regular drinking water or drinking water containing broad-spectrum antibiotic cocktail throughout the experimental period. **A)** One set of mice (n=8/groups) were tested for blood glucose levels, weekly for up to 12-weeks post-treatment initiation, to detect hyperglycemia/diabetes. **B)** One set of mice (5/group) were euthanized one-week post-treatment and H&E stained pancreatic sections were examined for insulitis as described for Fig. 5. Overall percentage of islets with different insulitis grades are shown. **C)** Single cell suspension of PnLN from mice euthanized one week post-treatment were stained for CD4+ cells with Foxp3 expression, and representative FACS graphs (left panel) and mean±SD values of 5 mice tested individually (right panel) are shown. Statistical significance was calculated by Log-rank test (for panel A), Fisher’s exact test (for panel B), or *t*-test (for panel C).

### Co-administration of YBG, Raldh substrate and β-cell-Ag results in expansion of Foxp3+ cells and better protection of NOD mice from T1D

YBG treatment appears to have a direct, in addition to microbiota-dependent, impact on enhancing gut immune regulation. Further, YBG-induced gut innate immune response involves increased Raldh1A2 and IL10 expression, and this regulatory innate response appears to contribute to protection of NOD mice from T1D. Therefore, whether YBG induced gut innate immune response can be exploited to promote β-cell-Ag specific immune regulation and better protection from T1D was determined. In one set of experiments, NOD-BDC-Foxp3-GFP mice were fed with YBG, BDC-peptide and/or retinol for a brief period (3 consecutive days and rested for 4 days) and examined for the frequencies of GFP+ (Foxp3+) T cells in the gut mucosa. As shown in Fig. 7, mice that received a combination of YBG, BDC-peptide and retinol showed a robust increase in Foxp3+ T cells in Peyer’s patch (PP) compared to other groups of mice. This brief treatment with YBG, BDC-peptide and Retinol combination also caused a significant increase in IL10 response in gut mucosa compared to mice that received individual agents (not shown).

**FIGURE 7:**
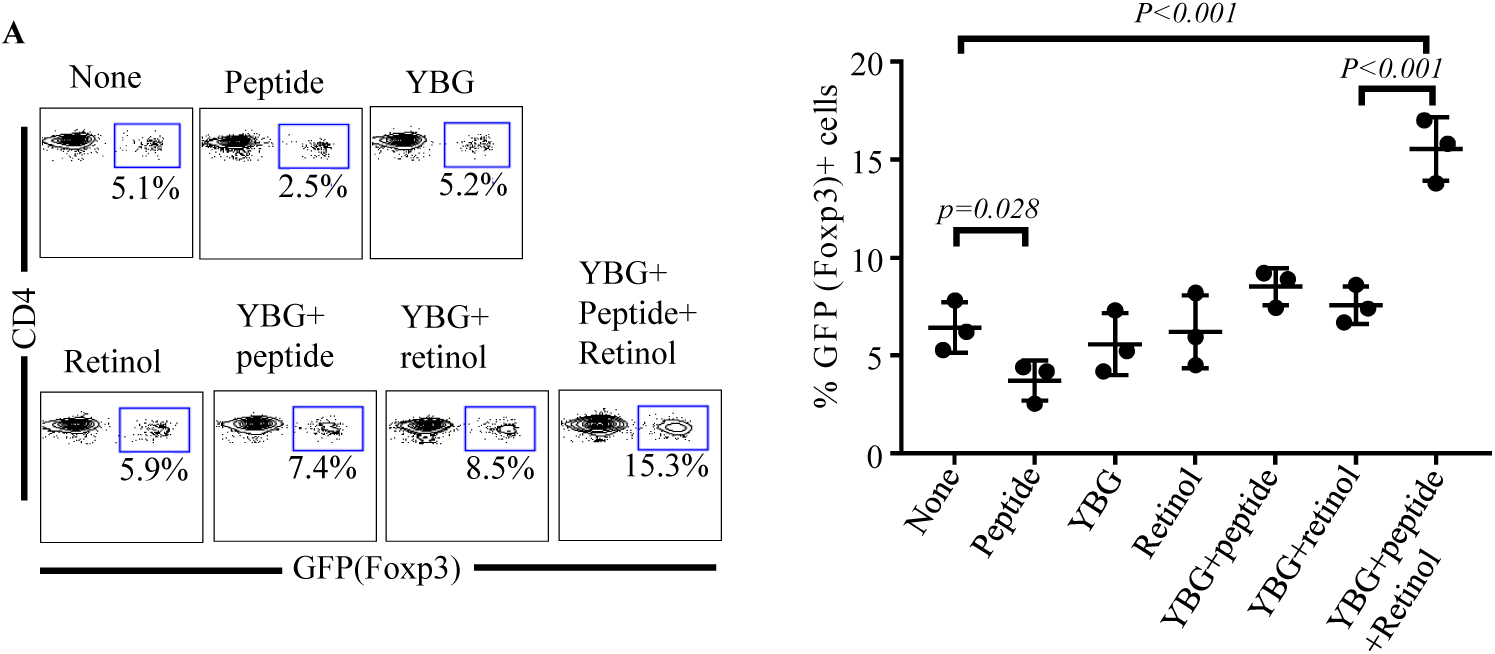
Short-term treatment with YBG, antigen and retinol resulted in a robust Treg response in vivo. Female NOD-BDC2.5-Foxp3-GFP mice were given YBG (250 μg/mouse/day), BDC2.5 peptide (5 μg/mouse/day) and/or retinol (0.5 μg/mouse/day) for 3 consecutive days, euthanized on day 7 and examined for GFP+(Foxp3+)CD4+ T cells in the Peyer’s patch by FACS. Representative FACS graphs (left panel) and mean±SD values of 3 mice/group tested individually (right panel) are shown. Statistical significance was calculated by *t*-test.

We, then, examined the impact of short-term co-administration of YBG, β-cell-Ag (peptide mix) and retinol on T1D in NOD mice. Ten to 12 week old pre-diabetic mice were treated with YBG, β-cell-Ag and retinol individually or in combination for 3 days, rested for 4 days, and the treatment was repeated for additional 3 days (a total of 6 days instead of 30 consecutive days of treatment as described for Fig. 4), and monitored for hyperglycemia or examined for insulitis and immune cell phenotypes. As shown in Fig. 8A, while this short-term treatment with YBG alone, or other individual agents, did not produce substantial protection from T1D, mice that received YBG, β-cell-Ag and retinol combination showed a significant delay in hyperglycemia compared to untreated controls as well as YBG treated mice. Pancreatic islets of mice that received this combination treatment showed less severe insulitis compared to that of control groups (Fig. 8B). Importantly, PnLN cells from mice that received this combination treatment showed relatively higher frequencies of Foxp3+ T cells (Fig. 8C). Furthermore, higher frequencies of IL10+ and IL17+ and lower frequencies of IFNγ+ cells were detected in ex vivo β-cell-Ag challenged PnLN from mice that received the combination therapy (Fig. 8D and Supplemental fig. 4). PnLN cells from these mice also secreted higher amounts of IL10 and IL17 and lower amounts of IFNγ upon ex vivo challenge with β-cell-Ag (Fig. 8E). T cells enriched from the pancreatic tissues of mice that received the combination treatment also showed a similar trend in Foxp3+, IFNγ+ and IL10+ Tregs supplemental fig. 5). Mice that received short-term treatment with YBG or retinol alone showed no profound difference in cytokine response. Of note, these pre-diabetic mice that received oral administration of β-cell-Ag alone showed not only severe insulitis and rapid progression to hyperglycemia, but also secreted higher pro-inflammatory IFNγ and lower IL17 and IL10 compared to untreated controls. Lower Foxp3+, IL10+ and IL17+ cell, and higher IFNγ+, cell frequencies were also detected in the PnLN of mice that received β-cell-Ag alone. Overall, these observations suggest that while oral administration of β-cell-Ag alone, at close to disease onset, could accelerate the disease progression by enhancing pro-inflammatory Th1 cytokine response and suppression of Treg function, YBG induced gut regulatory immune response can be exploited to enhance/promote oral tolerance against self-antigens, even at advanced stages of disease progression, to prevent T1D.

**FIGURE 8:**
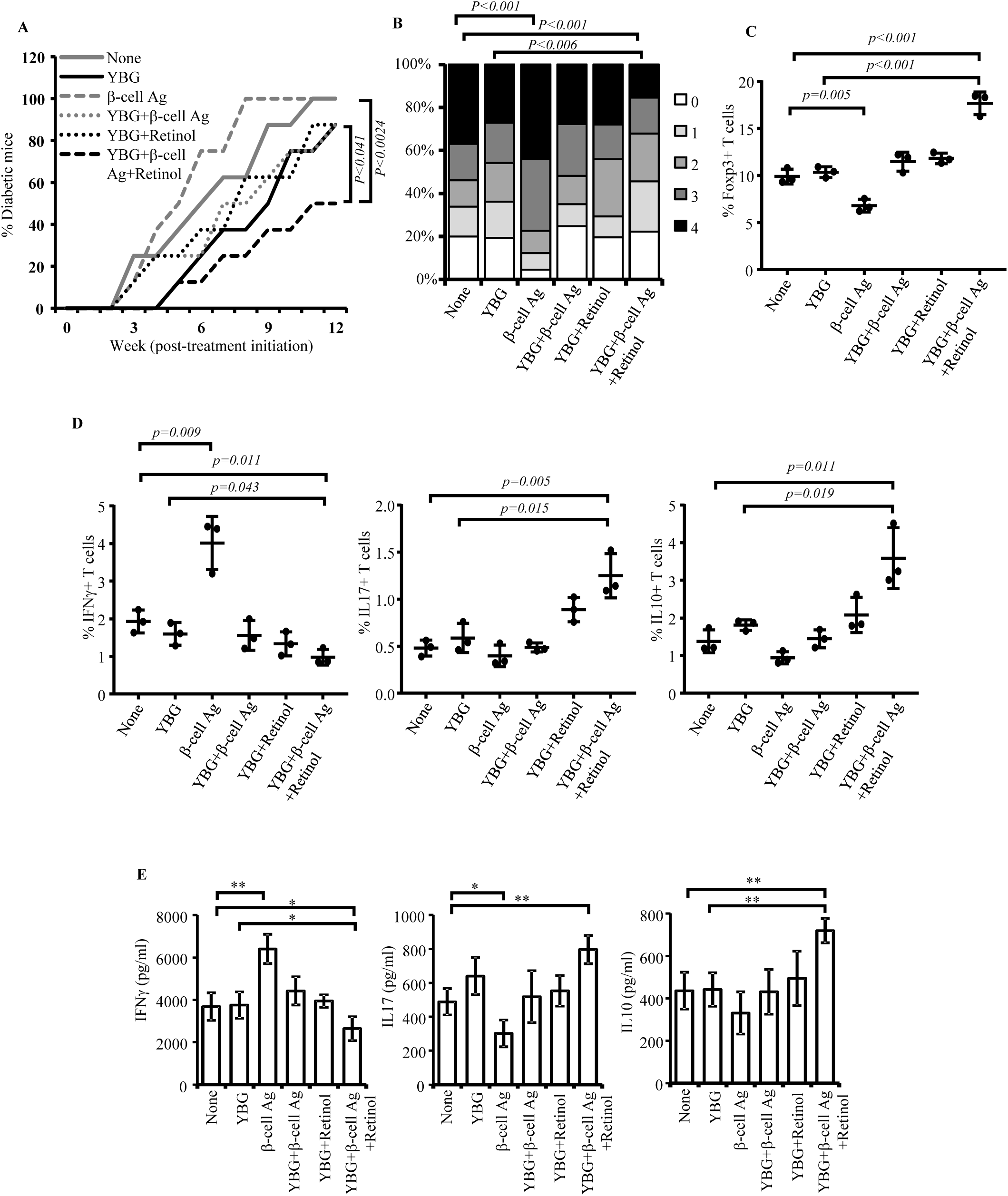
Short-term treatment with YBG, β-cell-Ag and retinol resulted in a robust modulation of immune response and better protection of NOD mice from T1D. 10-12-week old female NOD mice (8 mice/group) were given YBG, β-cell-Ag peptide mix and/or retinol as described above for panel A and the treatments were repeated once with a 4-day interval. **A)** One set of mice was tested for blood glucose levels weekly for up to 12-weeks post-treatment initiation and percentage of mice in each group with hyperglycemia are shown (left panel). **B)** One set of mice (4 mice/group) was euthanized, two-week post-treatment and H&E stained pancreatic sections were examined for insulitis. Overall percentage of islets with different insulitis grades are shown. **C)** Fresh PnLN cells from one set of mice (3 mice/group) euthanized 7 days post-treatment were examined for Foxp3+ T cell frequencies by FACS. **D)** PnLN cells from an additional set of mice were cultured with β-cell-Ag peptide mix for 24 h and examined for intracellular cytokine levels by FACS. Mean ±SD values of 3 mice/group tested individually are shown for panels C and D. Representative FACS graphs are shown in **Supplemental fig. 4**. **E)** PnLN cells of mice from a similar experiment were cultured with β-cell-Ag peptide mix for 48 h and supernatants were tested for cytokine levels by ELISA. Statistical significance was calculated by Log-rank test (for panel A), Fisher’s exact test (for panel B) or *t*-test (for panels C-E).

## Discussion

A large body of literature suggests that certain complex dietary polysaccharides such as BGs of different microbial and plant origin affect the host immune function ^27-29, 38, 39^. Since much of this literature consists of *in vitro* experiments or studies in which these polysaccharide preparations were systemically injected, the true impact of orally administering BGs on gut immune function is largely unknown. Also, use of poorly defined and crude polysaccharide preparations, BG of yeast origin in particular, has contributed much to confusion, misconceptions, and controversies regarding its biological effects. Hence, the true immunologic effects following systemic and oral administration of well-defined β-1,3/1,6-glucans and the impact on autoimmunity are largely unknown. In this regard, our recent report showed that systemic administration of low-dose purified YBG prevents autoimmune diabetes in NOD mice by inducing an immuno-adjuvant effect ^4^ suggesting that β-1,3/1,6-glucan interaction with innate immune receptors in the systemic compartment enhances overall immune regulation under autoimmune susceptibility. Since orally consumed dietary polysaccharides can influence the gut immune function by directly interacting with cell surface receptors and indirectly through gut microbes ^28, 29, 40^, in this study, we determined the impact of oral administration of purified YBG as a dietary supplement on gut immune function, fecal microbiota composition and autoimmune diabetes. Here, we show that oral administration of YBG results in enhanced gut immune regulation, altered gut microbiome, and protection of NOD mice from T1D. We also show that YBG not only has a pre-biotic effect on gut microbes, but also has a direct immune regulatory oral-adjuvant property, which can be exploited to modulate autoimmunity for preventing T1D.

Several reports have shown that the biological effect of β -1,3/1,6-glucans such as YBG is the result of interaction with specific receptors like Dectin-1 on immune cells ^4-6^. Importantly, it is now recognized that gut resident DCs have the ability to interact directly with dietary and microbial antigens of gut lumen ^30-32^. In fact, our experiments using WT and Dectin-1 deficient mice show that the overall and IDC specific immune response of small intestine to orally-administered purified YBG is primarily Dectin-1 dependent. Interestingly, as indicated by the ability of IDCs to promote Treg induction/expansion as well as to modulate the cytokine response of T cells, YBG induced gut innate immune response is immune regulatory in nature.

Colonic microbial communities can ferment non-digestible dietary polysaccharides and use them as energy sources for their growth ^41-43^. Therefore, these polysaccharides are being considered as prebiotics with the ability to promote growth of specific beneficial microbial communities. In fact, we observed that prolonged oral treatment using low-dose YBG causes changes in the gut microbial composition by reducing the abundance of Firmicutes and increasing the abundance of Bacteroidetes and Verrucomicrobia phyla, which include many polysaccharide fermenting bacteria. Further, predictive functional profiling of fecal microbiota shows overrepresentation of metabolic pathways linked to carbohydrate metabolism, glycan biosynthesis and metabolism, and biosynthesis of secondary metabolites after YBG treatment compared to pre-treatment time-point.

Previous studies have shown that gut colonization by specific microbial communities causes induction and expansion of Foxp3+ Tregs in the intestinal and systemic compartments ^44-46^. Further, gut bacteria play a critical role in maintaining peripheral tolerance and gut immune homeostasis ^47, 48^. Our results show that depletion of gut bacteria using broad-spectrum antibiotics diminishes Foxp3+ T cell frequency, compared to mice with intact gut microbiota, but less significantly in YBG treated mice compared to their control counterparts. Our observations that both control and bacteria-depleted mice, which received YBG, have higher frequencies of Tregs in the small intestine compared to their control counterparts with intact microbiota suggest that YBG-treatment associated increase in Treg abundance is, at least in part, the result of its direct interaction with gut mucosa. On the other hand, our results show that YBG treatment induced enhanced immune regulation in the large intestine is primarily microbiota-dependent. Nevertheless, future studies using gnotobiotic mice conventionalized using control and YBG-shaped microbiota will be needed to conclusively determine whether YBG treatment-shaped gut microbiota contributes to Treg induction/expansion and overall immune regulation.

Progression of autoimmune diseases including T1D in at-risk subjects and rodent models can be influenced by dietary factors and gut microbiota ^49-52^. Importantly, clinical trials and pre-clinical studies have exploited the tolerogenic nature of gut mucosa for inducing self-antigen specific immune modulation to prevent autoimmunity ^53-55^. Shaping the gut microbiota as well as enhancing gut immune regulation by employing dietary factors presents attractive approaches for modulating autoimmune diabetes and enhancing the efficacy of tolerogenic oral vaccines. Our observations that prolonged oral treatment with low-dose YBG modulates T cell function in the systemic compartment, suppresses insulitis progression and delays T1D incidence suggest that YBG treatment induced-regulatory innate immune response, in fact, has a direct suppressive effect on autoimmunity against pancreatic β-cells. Interestingly, we found that YBG treatment associated gut innate immune response involves higher expression of Raldh along with various anti-and pro-inflammatory cytokines. Raldh enzymes, which convert retinol to retinoic acid, are considered as the key tolerogenic factors expressed in gut resident DCs. Importantly, the ability of retinoic acid to promote the conversion of antigen recognized effector T cells to Tregs has been well recognized ^56-59^. Our observation that the oral co-administration of YBG, peptide antigen and retinol results in a robust increase in Tregs in the gut mucosa suggests that tolerogenic self-antigen presentation in the gut mucosa could be enhanced through YBG induced innate immune response. This notion has been substantiated by the observation that a better protection of NOD mice from insulitis and hyperglycemia can be achieved by oral co-administration of YBG, peptide antigen and retinol compared to administering individual agents.

Overall, our observations show that Dectin-1 dependent innate immune response induced by YBG in the gut mucosa involves both immune regulatory and pro-inflammatory factors. We show that, in addition to causing changes in the gut microbial composition which appears to have an impact on gut immune regulation, the pro-inflammatory and immune regulatory responses triggered by YBG in the gut mucosa can promote both Treg and Th17 responses, leading to suppressed Th1 dominated β-cell Ag specific autoimmunity of systemic compartment and protection of NOD mice from T1D. Others and we have shown that Th17 response independently or in combination with Treg response has a protective effect against β-cell autoimmunity in NOD mice ^4, 21, 60^. Most importantly, our study also shows that YBG induced regulatory innate immune response of the gut mucosa can be exploited to modulate T cell response to pancreatic β-cell-Ag, for achieving better protection from T1D, highlighting the oral tolerogenic adjuvant value of low-dose YBG treatment.

## Supporting information

## Author contribution

R.G. researched and analyzed data, and edited the manuscript; N.P. researched data and reviewed manuscript; B.J. researched data and reviewed manuscript; M.H.S. researched data and reviewed manuscript; S.Q. researched data and reviewed/edited manuscript; R.B. researched data and reviewed manuscript; S.K.M. researched data and reviewed manuscript; and C.V. designed experiments, researched and analyzed data, and wrote/reviewed/edited manuscript.

## Acknowledgments

This work was supported by internal funds from MUSC and UIC, National Institutes of Health (NIH) grants: R21AI133798 (NIAID) and R21AI133798 Administrate supplement (ODS), and American association of diabetes grant ADA-1-13-IN-57 to CV. The authors are thankful to the Histology core of Pathology department of MUSC for the histology service, and Genomic center of MUSC for 16S rRNA gene sequencing service. Dr. Vasu is the guarantor of this work and as such, has full access to all the data in the study and takes responsibility for integrity of the data and accuracy of data analysis.

