## Supplementary material for "Complex dietary-polysaccharide modulates gut immune function and microbiota, and promotes protection from autoimmune diabetes"

Supplemental fig. 1

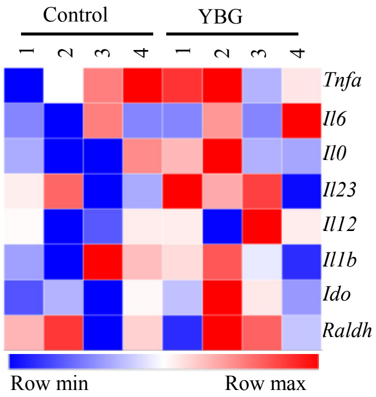


**Supplemental fig. 1: Effect of short-term oral administration of low dose YBG on small intestine**. Eight-week-old WT B6 mice were given YBG suspension (250 μg/mouse/day) by oral gavage for 3 consecutive days and euthanized <24 h post-treatment to harvest the intestine. cDNA prepared from the distal ileum region were subjected to qPCR and the expression levels of cytokines and non-cytokine factors were compared. Expression levels relative to β-actin expression were plotted as a heatmap with Morpheus application. n= 4 mice/group and the assay was performed in triplicate for each sample.

Supplemental fig. 2

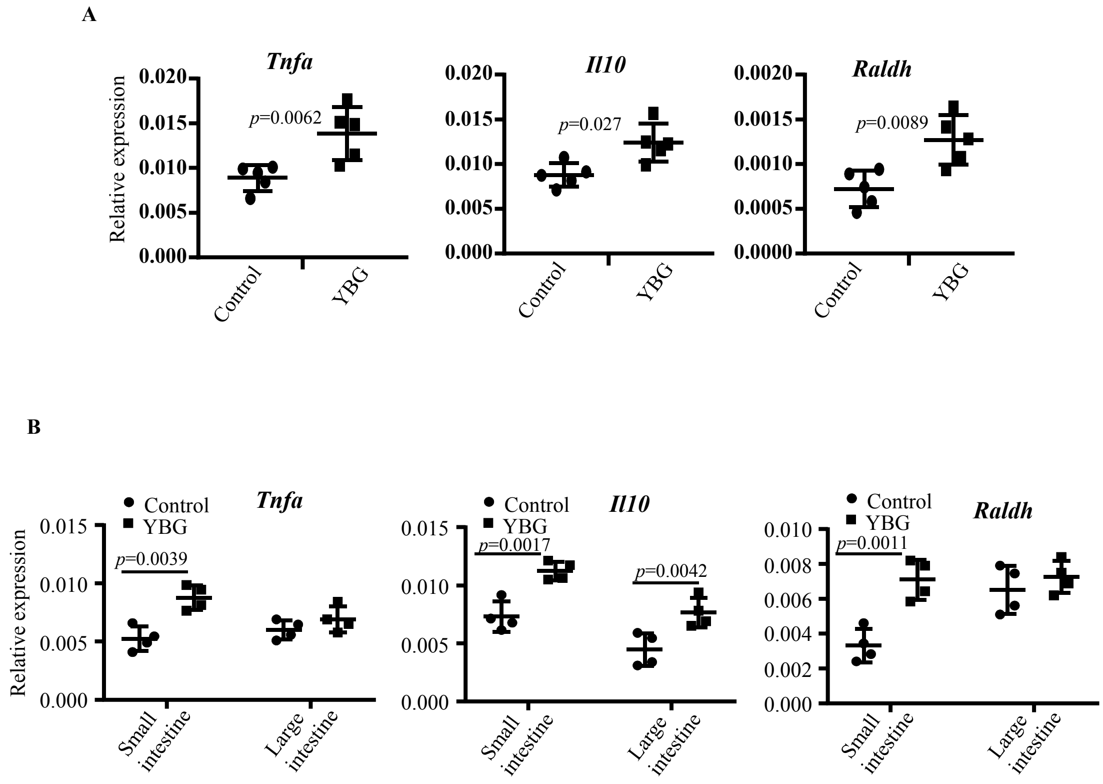


**Supplemental fig. 2: Effect of prolonged oral administration of low-dose YBG on intestine of NOD (A) and B6 (B) mice**. Ten week old NOD mice or 8 week old B6 mice were given YBG suspension (250 µg/mouse/day) by oral gavage for 30 consecutive days and euthanized 24 h post-treatment to harvest the intestine. cDNA prepared from the distal ileum (for panel A) or distal ileum and colon (for panel B) were subjected to qPCR and the expression levels of cytokines and non-cytokine factors were compared. Expression levels relative to β-actin expression were plotted. n= 5 mice/group for panel A and n=4 mice/group for panel B.

Supplemental fig. 3

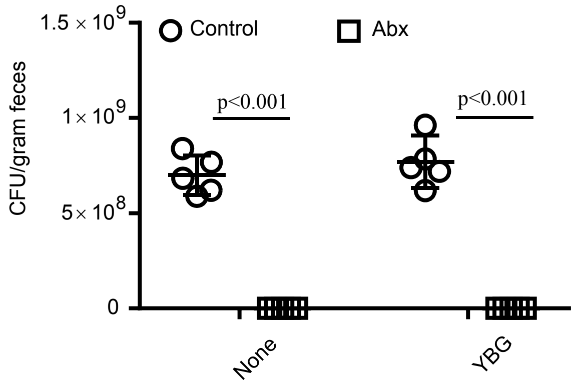


**Supplemental fig 3: Depletion of gut microbiota using antibiotics**. NOD mice were given a broad-spectrum antibiotic cocktail (ampicillin (1 g/l), vancomycin (0.5 g/l), neomycin (1 g/l), and metronidazole (1 g/l) - containing drinking water for up to 30 days and treated with YBG as described for Fig. 6. Fecal pellets collected from individual mice on day 30 were suspended and diluted in sterile PBS, plated on to brain heart infusion plates under anaerobic and aerobic conditions for up to 72 h, the total number of colonies were counted, and colony forming units (CFU)/gram initial fecal material were calculated. n= fecal pellets from 5 representative mice/group.

Supplemental fig. 4

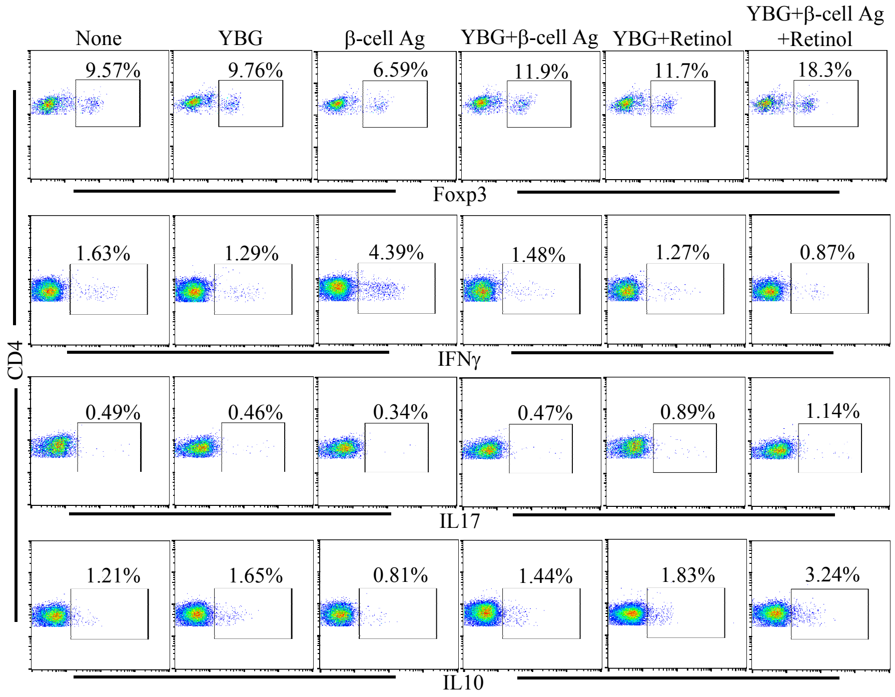


**Supplemental fig 4: Short-term treatment with YBG, β-cell-Ag and retinol resulted in a robust modulation of immune response**. NOD mice were treated as described for Fig.8 and fresh PnLN cells from one set of mice (3 mice/group) euthanized 7 days post-treatment were examined for Foxp3+ T cell frequencies by FACS. PnLN cells were also cultured with β-cell-Ag peptide mix for 24 h and examined for intracellular IFNγ, IL17, and IL10 by FACS. Representative FACS plots are shown here. Mean ±SD values are shown in Fig. 8.

Supplemental fig. 5

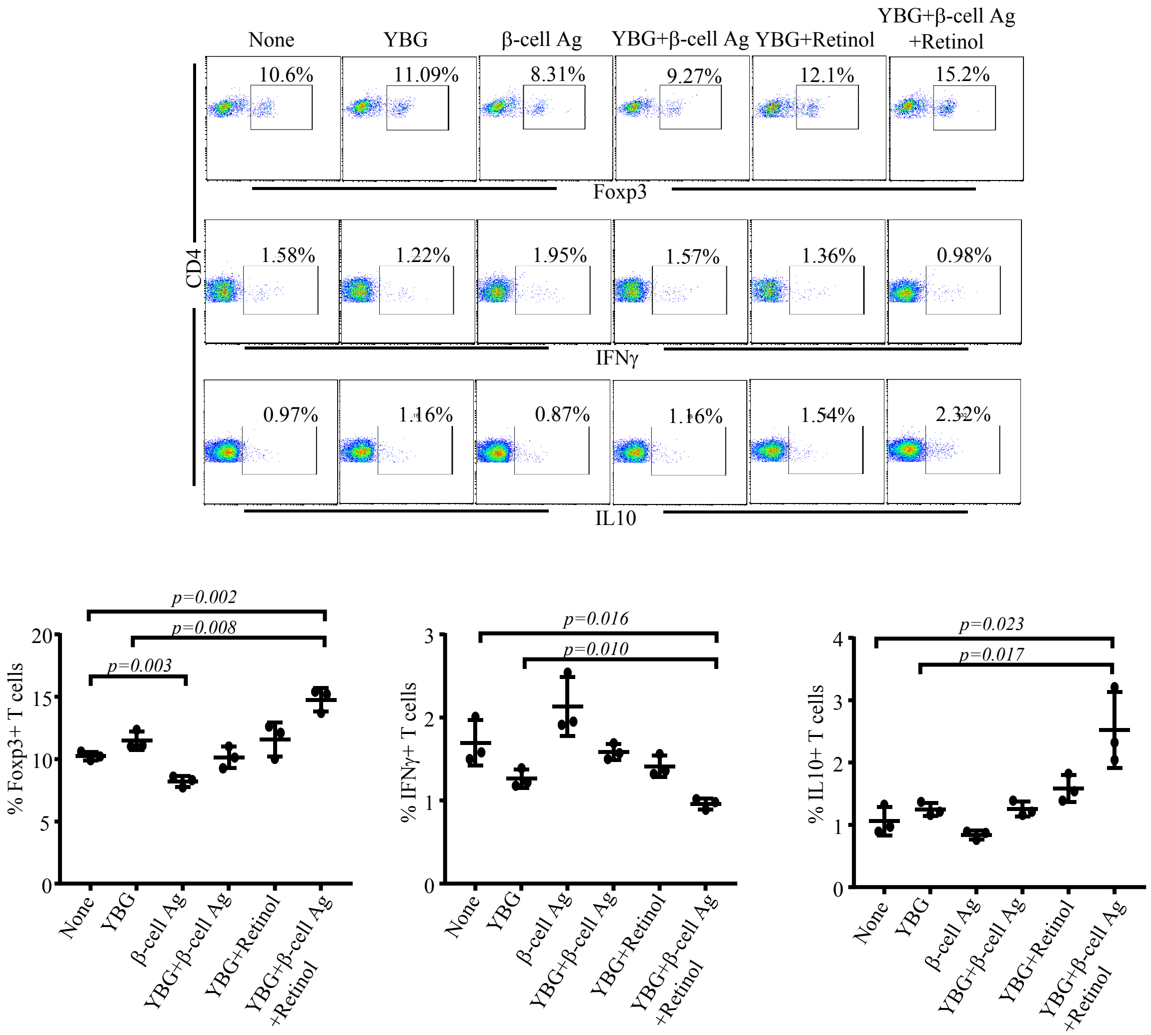


**Supplemental fig 5: Short-term treatment with YBG, β-cell-Ag and retinol resulted in a modulation of immune response**. Pre-diabetic NOD mice were treated as described for Fig.8 and immune cells were enriched from whole pancreas of one set of mice (3 mice/group) euthanized 7 days post-treatment by sequential collagenase digestion, trypsin dissociation, and Percoll (35% and 70%) gradient centrifugation. Immune cell rich fraction from 35%-70% Percoll interphase were examined for Foxp3+ T cell frequencies by FACS. Pancreatic immune cells were also cultured with β-cell-Ag peptide mix for 24 h and examined for intracellular IFNγ and IL10 by FACS. Representative FACS plots (upper panel) and mean ±SD values (lower panel) are shown
